# Recovering Single-cell Heterogeneity Through Information-based Dimensionality Reduction

**DOI:** 10.1101/2021.01.19.427303

**Authors:** Benjamin DeMeo, Bonnie Berger

**Affiliations:** Computer Science and Artificial Intelligence Laboratory, MIT, Cambridge, MA 02139, USA; Department of Biomedical Informatics, Harvard University, Cambridge, MA 02138, USA; Department of Mathematics, MIT, Cambridge, MA 02139, USA

## Abstract

Dimensionality reduction is crucial to summarizing the complex transcriptomic landscape of single cell datasets for downstream analyses. However, current dimensionality reduction approaches favor large cellular populations defined by many genes, at the expense of smaller and more subtly-defined populations. Here, we present surprisal component analysis (SCA), a technique that leverages the information-theoretic notion of *surprisal* for dimensionality reduction, and demonstrate its ability to improve the representation of clinically important populations that are indistinguishable using existing pipelines. For example, in cytotoxic T-cell data, SCA cleanly separates the gamma-delta and MAIT cell subpopulations, which are not detectable via PCA, ICA, scVI, or a wide array of specialized rare cell recovery tools. We also show that, when used instead of PCA, SCA improves downstream imputation to more accurately restore mRNA dropouts and recover important gene-gene relationships. SCA’s information-theoretic paradigm opens the door to more meaningful signal extraction, with broad applications to the study of complex biological tissues in health and disease.

## Introduction

Single-cell RNA sequencing (scRNA-seq) produces transcript counts for individual cells, enabling fine-grained analyses of biological tissues. In particular, single-cell datasets can uncover cellular populations that play critical roles in biological and pathological phenomena [1, 2], and identifying and characterizing these populations is a key motivator of many single-cell experiments.

However, the size, high dimensionality, and noisiness of single-cell data complicates this task. Modern experiments profile tens of thousands of genes per cell, often with high dropout levels and technical noise. Consequently, dimensionality reduction, whereby the data is represented in a lower-dimensional space with enriched signal, has become a cornerstone of modern scRNA-seq analysis pipelines. For example, Principal Component Analysis (PCA) projects the data to a lower-dimensional linear subspace such that the total variance of the projected data is maximized. Independent Component Analysis (ICA) instead aims to identify non-Gaussian combinations of features. Both have found widespread use in single-cell studies [3, 4, 5, 6]. A popular recent method, scVI [7], models transcript counts using a zero-inflated negative-binomial distribution, and performs variational inference to non-linearly embed each cell into a low-dimensional parameter space. scVI’s non-linearity supports more complex functions between a cell’s transcripts and its identity, but the dimensions of the latent space are harder to interpret than those of PCA and ICA.

While these approaches are effective, they often fail to capture the full cellular diversity of complex tissues for two reasons. First, rare cell types, by definition, account for a small fraction of the observations, and therefore contribute little to a dataset’s global structure. Second, many distinctions between cellular populations hinge on just a few of the thousands of genes measured. For instance, gamma-delta T-cells are distinguished from ordinary cytotoxic T-cells by the presence of just a few gamma and delta T-receptors; we call such populations *subtly-defined*. Whereas PCA and ICA both compute features that optimize objective functions over the entire dataset, i.e. total variance and non-Gaussianity, rare cell populations thwart both strategies since the genes defining them may be noisy or unexpressed over much of the data. Subtly-defined populations present an additional challenge, since the few discriminative genes may be masked by many irrelevant genes. Similarly, scVI uses the evidence lower bound (ELBO) loss function to evaluate and refine its latent encoding. Since ELBO takes each recorded transcript into account, rare and subtly-defined cell types may not impact it much, leading to under-representation. Real cellular populations may fall into one or both of these categories, and thus these challenges are significant roadblocks to realizing scRNA-seq’s full potential.

Here, we introduce SCA (surprisal component analysis), an information-theoretic dimensionality reduction method that identifies statistically informative signals in transcriptional data to enable deeper insight into complex tissues (Figure 1a). SCA leverages the notion of *surprisal*, whereby less probable events are more informative when they occur, to assign an *surprisal score* to each transcript in each cell. In doing so, SCA enables dimensionality reduction that better preserves information from rare and subtly-defined cell types, uncovering them where existing methods cannot.

**Figure 1:**
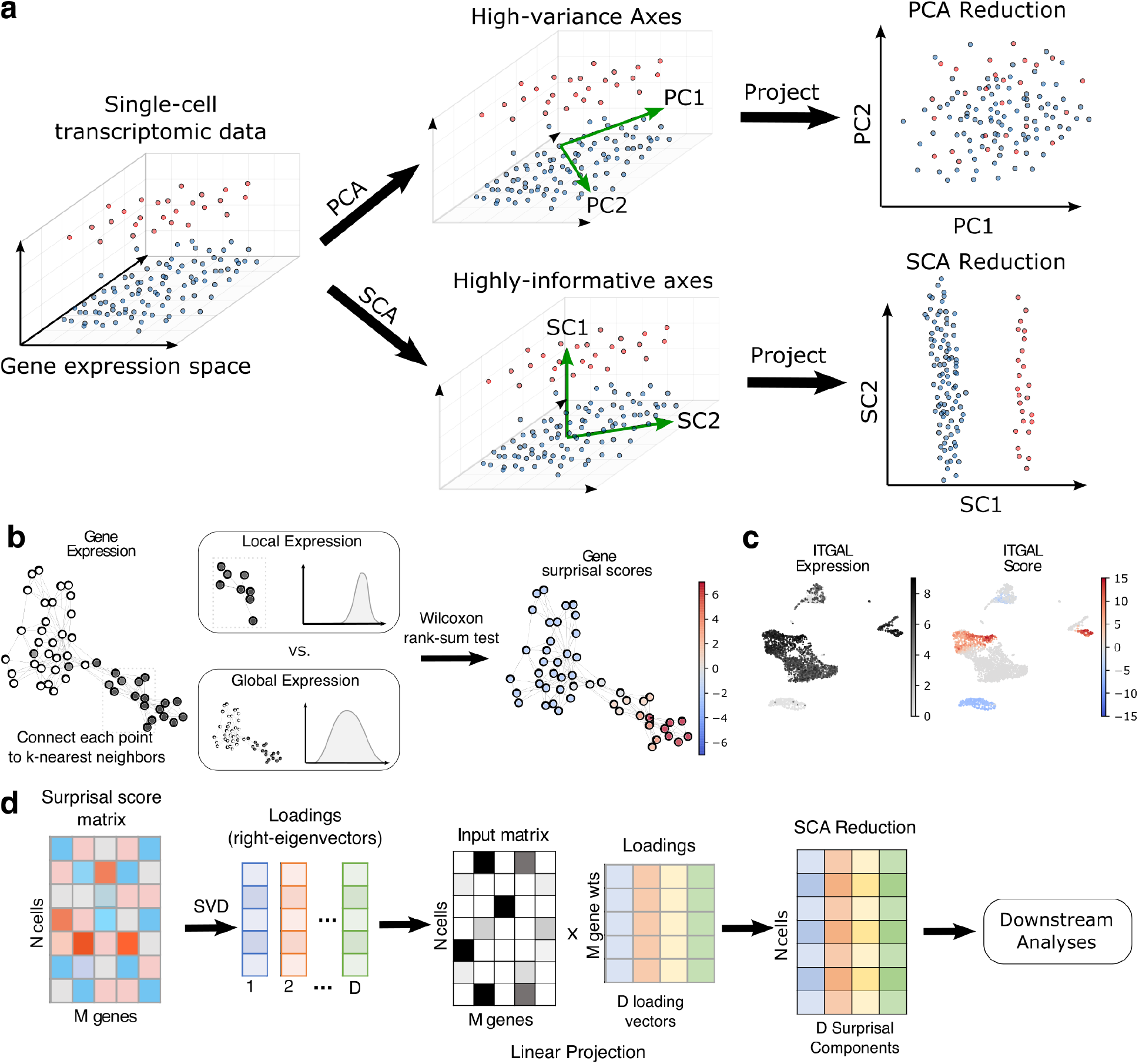
**a**: Illustration of SCA’s main conceptual advance. The vertical axis separates a small cellular population (top) from a larger one (bottom). The two horizontal axes have higher variance, but cannot separate the two populations. The leading Principal Components align with the higher-variance horizontal axes align with the higher-variance horizontal axes, and fail to separate the populations. The leading surprisal component aligns with the more informative vertical axis, allowing downstream separation. **b**: Construction of surprisal scores from gene expression data. For each cell, we compare the gene’s expression in a local neighborhood of the cell to the gene’s global expression using the Wilcoxon rank-sum test. The resulting *p*-values are negative log-transformed to give the surprisal of the observed over- or under-expression, and given a positive sign for over-expression and a negative sign for under-expression. **c:** Example surprisal scores of the *ITGAL* gene over a set of PBMCs profiled via Smart-seq 3 [19]. Scores are positive where the gene is locally enriched, near zero where it represents noise, and negative where it is conspicuously absent. **d:** Construction of surprisal components. Singular value decomposition of the surprisal scores over all genes yields *D* loading vectors capturing informative axes in the data. We then linearly project the input data to these axes, producing a *D*-dimensional representation of each cell for downstream analysis.

To demonstrate the utility of our approach, we ran SCA on real and simulated data with rare and subtly-defined cellular populations, and assessed our ability to recover these populations downstream. For comparison, we also tested PCA, ICA, scVI, and six rare cell type discovery tools: RaceID [8], GiniClust [9], CellSIUS [10], FiRE [11], GeoSketch [12], and Hopper [13]. We show that SCA enables detection of small populations, such as gamma-delta T-cells and Mucosal-associated Invariant T (MAIT) cells, which are invisible to existing pipelines and yet critical to the study of tumor immunology [1, 2]. At the same time, SCA reductions better capture larger-scale differences between more common cell types, enabling multi-resolution analysis without the need for re-clustering [14]. Beyond rare cell type recovery, we find that SCA more accurately recovers gene-gene relationships and restores dropouts when used as a basis for imputation via the state-of-the-art MAGIC [15].

SCA is broadly applicable, requiring no information aside from the transcript counts, and generalizes to data composed of discrete cell types or continuous trajectories. The output components have a clear linear relationship with the original transcripts, facilitating straightforward biological verification and interpretation. More broadly, we believe that SCA’s information-theoretic approach is a mathematically justified and empirically useful approach to signal extraction in any high-dimensional data modality, biological or otherwise.

## Results

### Overview of SCA

Like PCA and ICA, SCA projects the input data to a linear subspace spanned by a set of basis vectors, which we call *surprisal components*. SCA’s key conceptual advance is its novel approach to finding *informative* axes of variation, where an informative axis is one that separates cell types or captures biologically meaningful variation (Figure 1a). Past single-cell experiments have shown that the presence or absence of a small number of genes can determine a cell’s phenotype [5, 16, 17]. The key challenge, then, is to find and isolate these signals for each cell.

To this end, SCA first quantifies the importance of each transcript in each cell by converting transcript counts into *surprisal scores* (Figure 1b, Algorithm 1). To determine the score of a given transcript in a given cell, we compare its expression distribution among the cell’s *k* nearest neighbors to its global expression, i.e. to the expected distribution of the transcript among a set of *k* cells randomly chosen from the entire dataset. A transcript whose local expression deviates strongly from its global expression is more likely to inform the cell’s location in relation to other cells, and therefore its identity. We quantify this deviation through a Wilcoxon rank-sum test, which produces a *p*-value representing the probability of the observed deviation in a random set of *k* cells. Following Shannon’s definition [18], the *surprisal* or *self-information* of the observed deviation is then defined as the negative logarithm of this its probability, i.e. as *-* log(*p*). This is a positive number which measures how surprising the transcript’s local expression is, in units of nats when the logarithm is natural (changing the base only scales the scores by a constant factor, which does not affect SCA’s output). To distinguish over-from under-expression, we flip the sign for under-expressed transcripts (Methods). The resulting scores are compiled into an *surprisal score matrix* with the same dimensionality as the input data.

This strategy gives genes high positive scores where they are markers (genes that distinguish a cell type from the rest), scores near zero where they represent noise, and low negative scores where they are conspicuously absent (Figure 1c). For example, consider a marker gene for a rare population. The gene is unexpressed over much of the data, but highly expressed in cells belonging to the population. Thus, for these rare cells, the local expression is far higher than would be expected by chance, so the gene receives a high score for these cells. Likewise, a gene expressed everywhere *except* on a rare population receives low scores on members of the population. On the other hand, a noisy gene with no bearing on cellular identity receives low scores everywhere, since its distribution on *k*-neighborhoods resembles that of random sets of *k* cells.

We next seek to distill the signal captured by the surprisal score matrix into a low-dimensional representation (Figure 1d, Algorithm 2). As shown in the Methods and in Supplementary Note 1, the right-eigenvectors of the surprisal score matrix represent highly informative linear combinations of genes, which we call *surprisal components* (SCs). The first *D* right-eigenvectors, which we denote *v*_1_, …, *v*_*D*_, then span a linear subspace onto which we project the input matrix *X*. The resulting *N × D* matrix is the output of SCA. We emphasize that while the construction of the surprisal score matrix and of *v*_1_, …, *v*_*D*_ is nonlinear, SCA’s output is a linear projection of its input to their span. This places SCA in the category of linear dimensionality reduction methods, together with PCA and ICA. In particular, this means that each of the output features is a weighted sum of the input features, enabling straightforward interpretation.

In summary, SCA accepts a transcript matrix and sets of *k*-nearest neighbors for each cell, finds transcripts that inform each cell’s locality, and distills this information into a smaller set of features. This process amplifies the locality signal of the input *k*-nearest neighbor data. For example, even if only 10% of a cell’s neighbors belong to the same population, this is still highly significant when the population comprises only 1% of the sample. SCA’s surprisal scores would reflect this, and the resulting components would better separate the rare cells, as we verify. The input neighborhoods can be specified arbitrarily, but by default SCA computes them via Euclidean distance on a PCA representation.

This signal-boosting step can be repeated: from an initial SCA reduction, we can compute *k*-neighborhoods using the Euclidean metric, use these neighborhoods to compute a surprisal score matrix, and perform singular value decomposition to compute another SCA reduction. As we show in both real and simulated data, this often improves the representation of rare and subtly-defined cell types (Figure 2b and 3f). Intuitively, we begin with a weak notion of locality (provided by PCA) and continually refine it. In our experiments, we have found that performance usually stabilizes after 3-5 iterations and remains stable thereafter (Figure 3f, Supplementary Figure S2). Note that regardless of the number of iterations, SCA’s output remains a linear projection of its input – iteration simply refines the subspace onto which SCA projects.

**Figure 2:**
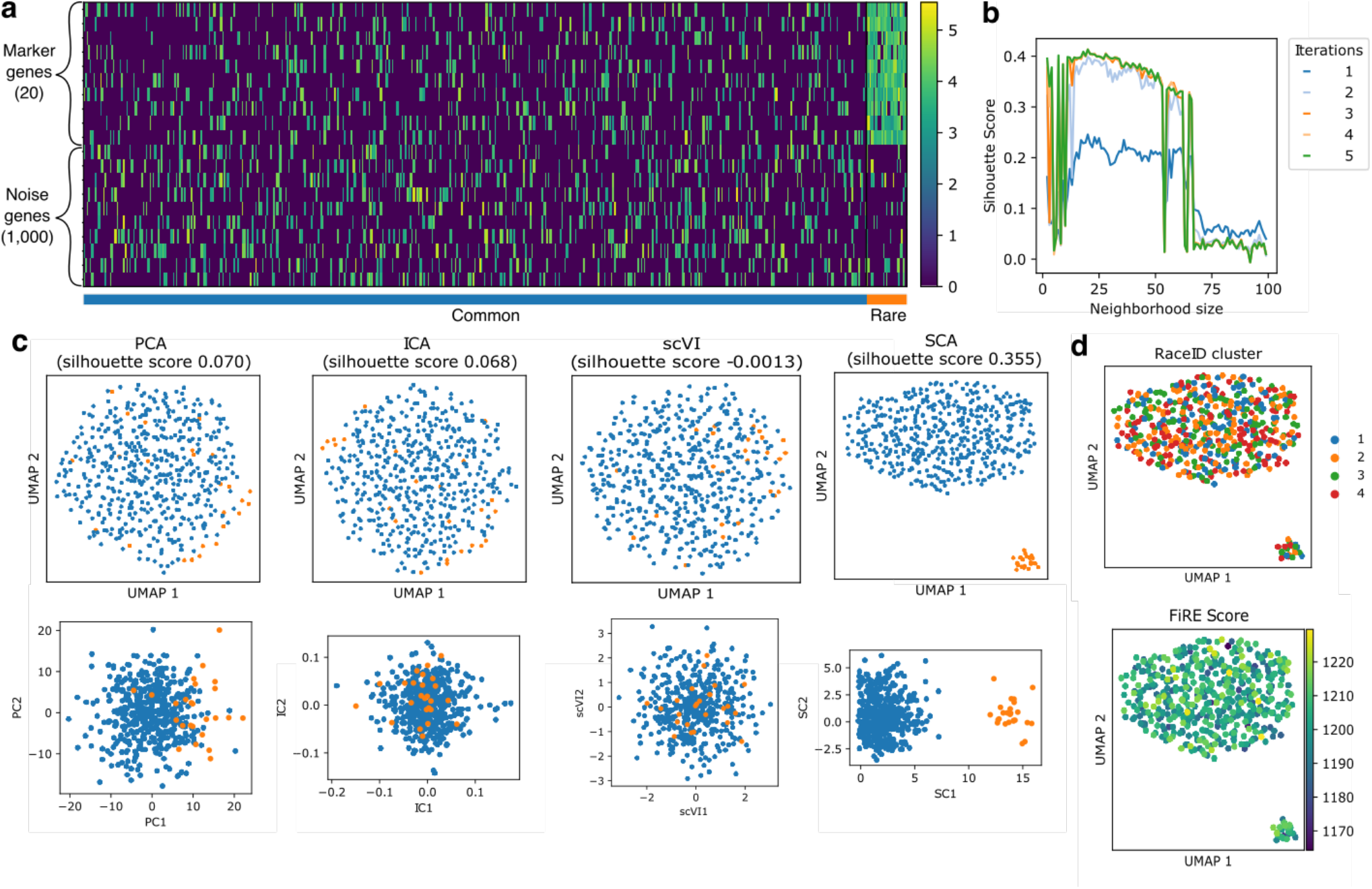
**a:** Illustration of synthetic dataset with 1,020 genes and 500 cells. 20 “marker genes” mark a set of 25 “rare” cells with a 10% dropout rate. The remaining 1,000 “noise” genes are expressed in a random 20% of the cells. To add further noise, each marker gene is also expressed in a random 15% of the remaining cells. Nonzero gene expression values are drawn randomly from the nonzero log-TPM gene expression values of a cytotoxic T-cell dataset profiled using Smart-seq 3 [19]. **b:** Silhouette scores for the two populations embedded via 20-dimensional SCA reductions computed with various neighborhood sizes and numbers of iterations. Performance is stable across neighborhood sizes up to 60, and improves with more iterations. **c:** Scatter plots of UMAP coordinates (top) and leading components (bottom) for PCA, ICA, scVI, and SCA (with 5 iterations). Top PCs, ICs, and scVI latent embedding dimensions do not separate the two populations. On the other hand, the leading surprisal component (SC) strongly separates the two populations, yielding complete separation in the downstream UMAP plot and a higher silhouette score. **d:** Performance of RaceID (top) and FiRE (bottom) on this dataset. RaceID clusters do not separate the populations, and FiRE scores are not higher for the rare cells.

**Figure 3:**
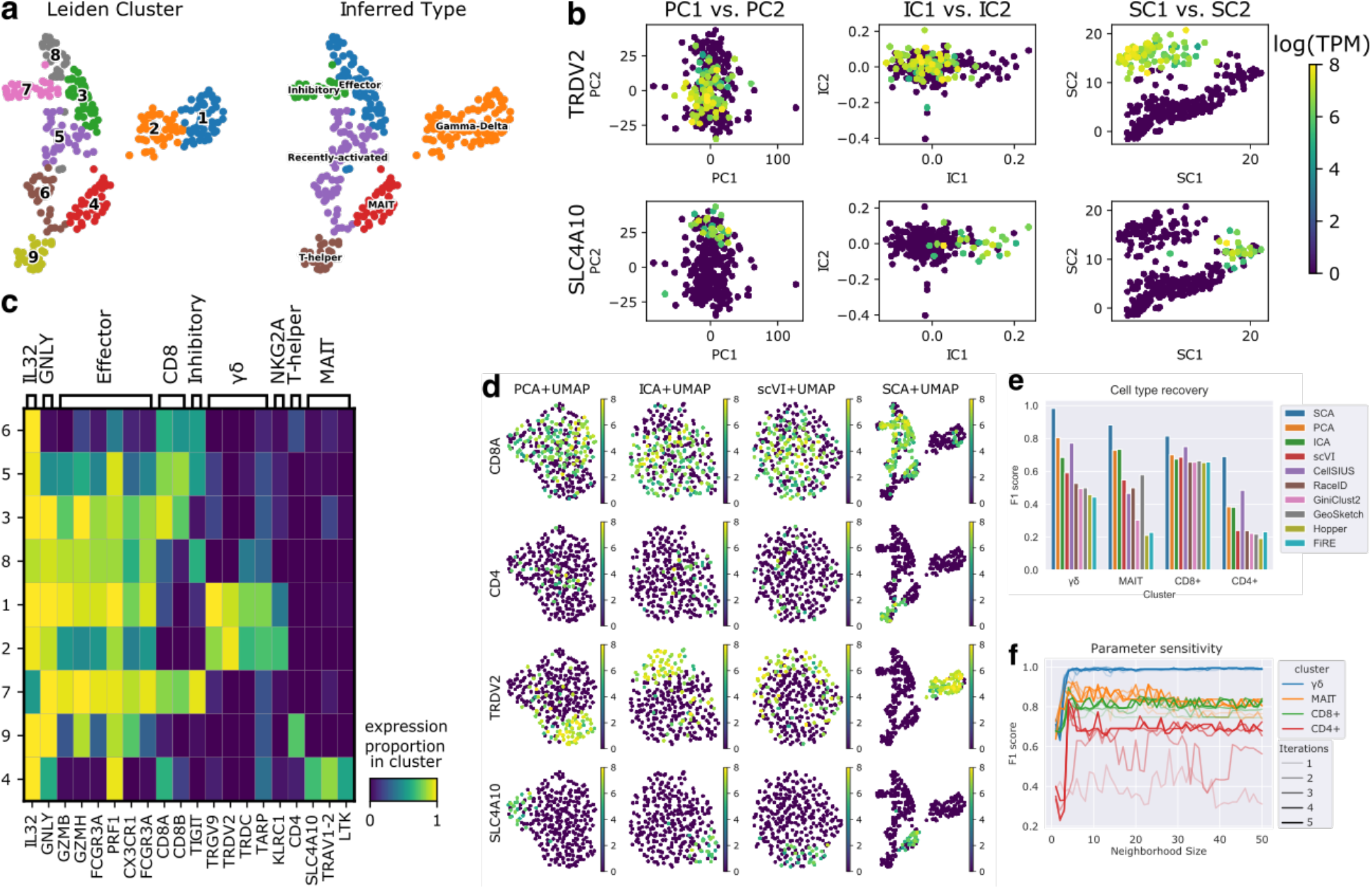
SCA recovers fine-grained cellular populations from a set of 307 cytotoxic T-cells profiled using Smart-seq 3 [19]. **a:** UMAP embedding computed from a 20-dimensional SCA representation using Euclidean nearest neighbors, with Leiden clusterings (left) and inferred cell types (right). Gamma-delta, MAIT, and T-helper populations cleanly separate. **b:** Scatter plots of leading Principal, Independent, and surprisal components, colored by log-TPM expression of key marker genes: the delta-receptor TRDV2 marks gamma-delta T cells [23], and SLC4A10 marks MAIT cells [24]. The leading surprisal components cleanly separate the gamma-delta and MAIT subpopulations, whereas the leading PCs and ICs blur these distinctions. **c:** Heatmap of gene expression percentages of key marker gene groups in each SCA-derived Leiden cluster. Gamma-delta, MAIT, and T-helper cells are clearly discernible. **d:** UMAP plots derived from 20-dimensional PCA, ICA, scVI, and SCA representations of the data, all using Euclidean nearest neighbors, colored by expression of key marker genes. CD8 T-cells, CD4+ T-helper cells, TRDV2+ gamma-delta T-cells, and SLC4A10+ MAIT cells form distinct regions of the SCA-derived UMAP plot, even though the other embeddings do not separate them. **e:** F1 scores for recovery of major T-cell populations by various clustering schemes (Methods). For PCA, ICA, and SCA, we assess Leiden clusters from the Euclidean 15-nearest neighbors graph with resolution 1. Leiden clusterings computed on the SCA representation consistently capture these cell types with superior accuracy. **f:** Robustness analysis for cell type recovery with respect to the size of the neighborhoods used to compute SCA’s surprisal scores, and the number of iteratrions. Performance improves with more iterations, and is stable across a wide range of neighborhood sizes.

### SCA recovers rare populations from noisy synthetic data

To test SCA’s power to recover rare cell populations in the presence of noise, we generated a synthetic dataset with 500 cells and 1,020 genes (Figure 2a). The genes are split into 20 “marker” genes and 1,000 “noise” genes, and the cells are split into 25 “common” cells and 475 “rare” cells. Each marker gene is expressed with 90% frequency in the rare cells, and 15% frequency in the common cells. Each noise gene is expressed uniformly at random throughout the data with frequency 20%. The nonzero expression levels are chosen at random from the nonzero log-TPM (transcripts per million) values of the cytotoxic T-cell dataset analyzed in Figure 3 [19], mimicking the erratic capture efficiency of real single-cell technologies. This dataset was designed to mimic a challenging biological scenario with a rare and subtly defined population and unreliable marker genes.

We reduced this dataset using 20-dimensional PCA; 20-dimensional ICA; scVI with a 20-dimensional latent space; and 20-dimensional SCA with 1-5 iterations (Methods). For each reduction, we assessed separation visually using UMAP coordinates [20], and quantitatively by assessing the silhouette score [21]. We found that the PCA, ICA and scVI representations collapsed the two populations, as indicated by very low silhouette scores (*<* 0.1) and poor separation in UMAP plots produced downstream of the Euclidean 15-nearest neighbors graph (Figure 2c). SCA produces increasingly better separations with successive iterations, and after 5 iterations achieves a silhouette score of 0.355, with a cleanly visible separation in the downstream UMAP plot (Figures 2b and 2c). This improvement is due to SCA correctly identifying the marker genes as having high information, and thus giving them higher impact on the final reduction; indeed, the first surprisal component clearly separates the two clusters, whereas the leading Principal and Independent components do not (Figure 2c, bottom). SCA’s performance is stable across a wide range of sizes of the *k*-nearest neighborhoods used to compute the surprisal score matrix, with a drop in performance only when the neighborhood size greatly exceeds the size of the rare population (Figure 2b).

For further comparison, we also tested four methods specifically designed for rare cell type recovery: GiniClust3 [9], CellSIUS [10], RaceID [8] and FiRE [11]. Neither RaceID’s clustering nor FiRE’s rarity scores separated the two clusters (Figure 2d). GiniClust3 did not identify any of the marker genes as having significantly high Gini index; consequently, it combined all cells into a single cluster. Similarly, CellSIUS did not identify any genes with bimodal distribution, and did not generate a clustering.

### SCA reveals the landscape of cytotoxic T-cell subtypes

Novel therapies increasingly leverage the immune system to fight disease, and the complexity and cellular diversity of immunological tissues makes them ideal targets for scRNA-seq [22, 5, 17]. However, these tissues also challenge the technology in a variety of ways: they contain diverse cell types with rare but clinically important sub-types, and the expression of individual surface receptors has outsize effects on phenotype [17, 22]. We therefore examined whether SCA can find and distill these signals to reveal a richer landscape of immune cell types.

To this end, we obtained a collection of 307 cytotoxic T-cells profiled using Smart-seq 3 (SS3) [19], and computed 20-dimensional reductions using PCA, ICA, and SCA. For SCA, we perform up to 5 iterations with a neighborhood size of 15. We followed each reduction with standard downstream steps: 15-nearest neighbor graph construction using the Euclidean metric, UMAP embedding, and Leiden clustering.

The PCA and SCA reductions both suggest that the data is homogeneous: the UMAP visualizations are globular, with no distinct clusters, and there is no obvious structure in the leading components (Figure 3b and 3d). On the other hand, SCA’s embedding reveals several clearly-separated populations, summarized by 9 Leiden clusters (Figure 3a). Differential gene expression analysis shows that these correspond neatly to well-documented biological cell types (Figure 1c). For example, clusters 1 and 2 contain Gamma-delta T-cells, as indicated by expression of the gamma- and delta-receptors *TRGV9* and *TRDV2* [23]. Cluster 4 uniquely expresses *SLC4A10, TRAV1-2*, and *LTK*, strongly suggesting MAIT cells [24, 2]. CD4+ T-helper cells group neatly in cluster 9, whereas high *TIGIT* levels in Cluster 7 suggest an inhibitory phenotype [25, 26]. Clusters 3 and 8 express standard cytotoxic effector genes like granzymes and perforins, whereas clusters 5 and 6 express CD8 but have low granzyme expression, suggesting recently-activated CD8 T-cells.

As shown in Figure 3d, the UMAP plots derived from the PCA and ICA representations do not separate cells based on key immunological markers, whereas the SCA-derived UMAP plot does. To quantify this and see how it affects *de novo* population discovery, we assessed whether the Leiden clusters computed downstream of each representation were concordant with a marker-based classification. We defined CD8+ T-cells as those expressing *CD8A*; CD4+ T-cells as those expressing *CD4*, gamma-delta T-cells as those expressing at least two of *TRGV9, TRDV2*, and *TRDC* ; and MAIT cells as those expressing at least two of *SLC4A10, TRAV1-2*, and *LTK*. For each Leiden clustering, we identified the set of clusters whose union is most concordant with each marker-based class, and computed the F1 score (Methods). For further comparison, we do the same for clusterings output by CellSIUS, RaceID, GiniClust, GeoSketch, and Hopper. To produce clusterings using Hopper, we perform Leiden-clustering on a 50-point Hopper sketch of the data, then assign each cell the cluster label of its nearest subsampled cell. For GeoSketch, we project the data to 20-dimensional PCA coordinates computed from a 50-point sketch, and then compute Leiden clusters. For FiRE, which assigns a continuous rarity score to each cell, we consider sets containing the top *k* highest-scoring cells for all values of *k* ranging from 1 to the full size of the dataset, and report the highest F1 score obtained by any such set. SCA consistently outperforms the other methods in identifying these immunological classes (Figure 3e).

For completeness, we performed robustness analysis to see how SCA’s performance varies under different choices of neighborhood size *k*, and different numbers of iterations (Figure 3f). As in the synthetic dataset, running more iterations often improves the representation; notably, the CD4+ sub-type is not consistently well-captured until at least the third iteration. After 3, 4, or 5 iterations, SCA performs well on all sub-types over a wide range of neighborhood sizes. Very small neighborhood sizes (*<* 5) do not perform as well, likely due to a lack of statistical power. Based on this and the synthetic data performance, we suggest choosing a neighborhood size small enough to be contained in the rarest population, but large enough to guarantee statistical power. We have found the default size of 15 to be suitable for many datasets.

### SCA distinguishes known cell types profiled by CITE-seq

We next sought to test whether SCA’s reductions better detect known populations of cells in a larger-scale immmunological single-cell dataset. We obtained a CITE-seq dataset from Hao et al. [5], in which hundreds of thousands of PBMCs from 8 human donors were subjected in parallel to transcriptomic profiling and to surface receptor profiling with a panel of 228 antibodies. The authors use both modalities, and input from human experts, to produce a cell type classification, which we take as a ground truth. Notably, the classification includes some T-cell populations which the authors were not able to detect through analysis of the transcriptomic data alone, such as gamma-delta and MAIT cells, offering a good challenge for SCA. We subset the data to T-cells, yielding 73,000 cells across all donors. To circumvent batch effects, we separate the data by donor and assess performance in each donor individually.

UMAP plots computed from each representation are shown for Patient 1 in Figure 4a, and similar UMAP plots for the remaining patients are provided in Supplementary Figure S4. Compared to PCA and ICA, SCA consistently generates more structured UMAP plots downstream, with clearer visual separation between cell types. We observe similar improvements when plotting the first two components of each reduction against each other (Figure S5). We hypothesized that this separation leads to more accurate clusterings downstream. To test this, we performed Leiden clustering [27] on each representation with resolution 1.0, and computed the adjusted mutual information (AMI) between the Leiden clusters and the true cell labels [28]. For comparison, we also computed AMIs for Leiden clusterings computed from scVI’s embedding, clusterings based on both GeoSketch and Hopper reductions, and clusterings returned by GiniClust3 [9], CellSIUS [10], and RaceID [8]. Across all patients, the SCA-derived Leiden clusterings achieved the highest AMI with the true cell labels (Figure 4b). Thus, SCA representations enable more accurate downstream recovery of cellular populations in a sample.

**Figure 4:**
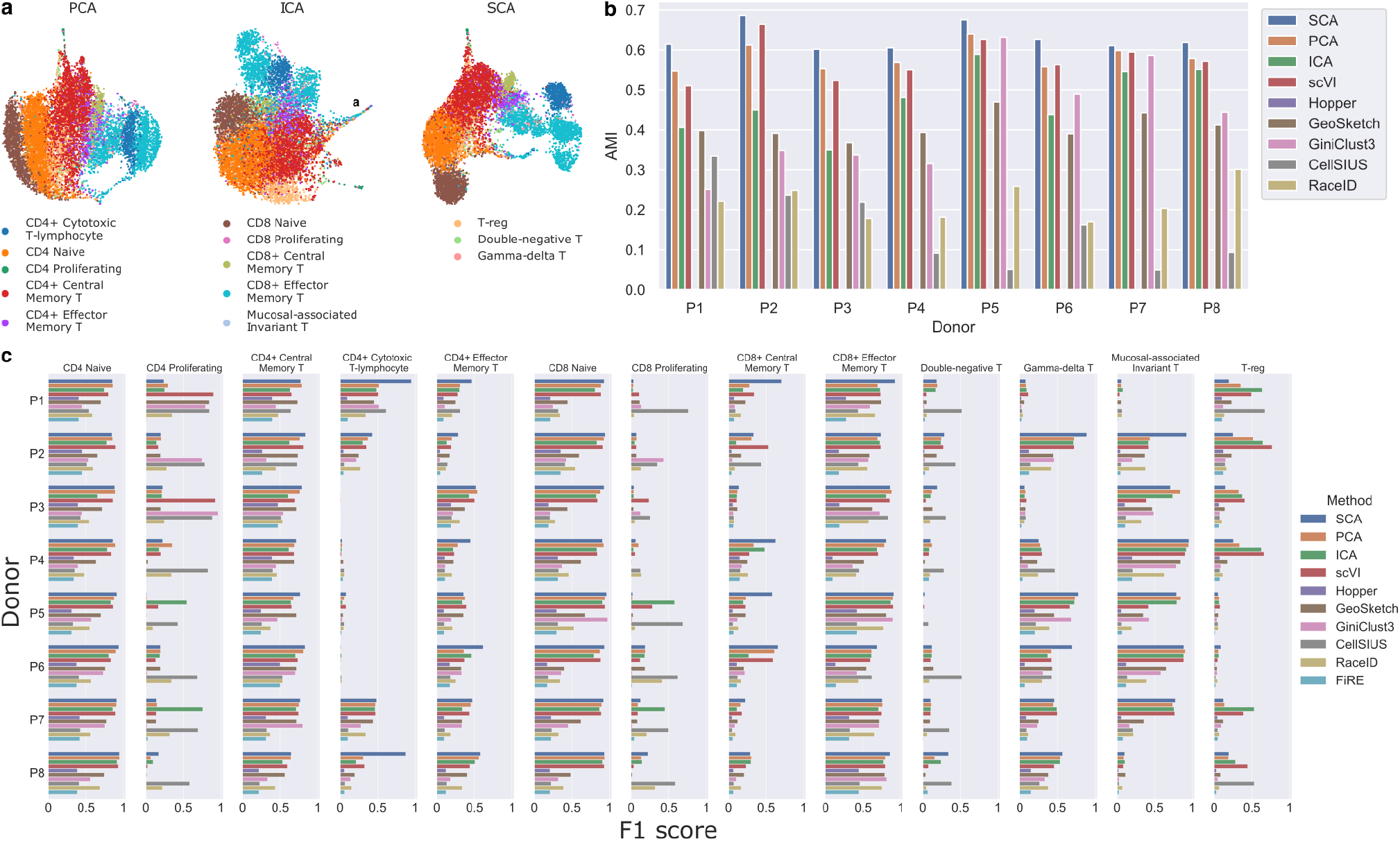
**a**: UMAP plots computed from PCA, ICA, and SCA reductions of T-cell scRNA-seq data for Patient 1 from Hao et al. [5]. Cell type labels were determined via parallel screening of 228 antibodies using CITE-seq. SCA produces better separation between the annotated cell types, enabling their discovery from transcriptomic data alone. **b**: Adjusted Mutual Information (AMI) of true cell labels with clusters output by each of the nine methods tested in each patient. (FiRE does not output a clustering but a rareness score, and thus is not amenable to AMI analysis.) For PCA, ICA, SCA, and scVI, we perform Leiden clustering with resolution 1.0 after reduction. SCA-based clusterings consistently have higher AMI with the true labels. **c**: F1 scores for recovery of all T-cell subtypes across all 8 patients of the dataset from Hao et al., from PCA, ICA, and SCA followed by Leiden clustering with resolution 1.0, and from seven other methods. For each clustering and cell type, the set of clusters best identifying that cell type was selected, and the F1 score reported. SCA recovers nearly all cell types with superior accuracy, and newly enables discovery of some types (e.g. CD4 cytotoxic T-lymphocytes and MAIT cells) which are poorly distinguished by other methods. Thus, SCA facilitates *de novo* discovery of these clinically relevant subpopulations.

We next assessed the accuracy with which SCA and other methods detect each of the known cellular populations, using the F1 score measure described above and in Methods. Across all donors, we find that clusterings derived from SCA recover most cell types with superior accuracy, with many cell types recoverable only by SCA (Figure 4c). For example, in Patient 1, SCA recovers CD4 cytotoxic T-lymphocytes (CD4 CTL) with F1 score 0.97, whereas no other method achieves a score higher than 0.61. We observe similar improvements across all patients, with SCA often recovering rare cell types (e.g. MAIT cells, Gamma-delta T-cells, and CD4+ Effector Memory T-cells) where existing methods do not. As notable exceptions, CellSIUS and GiniClust perform exceptionally well on the CD4 proliferating population, with CellSIUS also outperforming other methods on the CD8 proliferating and double-negative T populations. We suspect that the Leiden clusterings under-perform here due to theoretical limitations of community-detection algorithms to detect very small populations, as discussed in Kumpula et al. [29].

### SCA improves graph-based imputation

Dropouts and technical noise often obscure gene-gene relationships in single-cell data. Imputation aims to recover lost transcripts and restore these relationships. The widely-used MAGIC [15] tackles imputation by constructing a diffusion operator to share information across similar cells. By default, MAGIC computes cellular similarity using the Euclidean distance in PCA space. Since SCA better separates biological cell types, we pursued the intuition that using Euclidean distance in an SCA reduction would allow MAGIC to build a better diffusion operator resulting in more accurate imputation. We therefore newly formulated **SCA-MAGIC**, which performs diffusion over an SCA embedding instead of a PCA embedding (Methods).

To test the ability of each method to recover dropouts from the 500-cell synthetic dataset, we performed imputation using MAGIC and SCA-MAGIC, and measured the Pearson correlation between the imputed values and a ground-truth marker gene with no dropouts. The marker genes identifying the rare cell type population for the synthetic dataset each have a dropout rate of 10%. As shown in Figure 5a, SCA-MAGIC achieves significantly higher correlation between imputed marker genes and the ground-truth marker gene.

**Figure 5:**
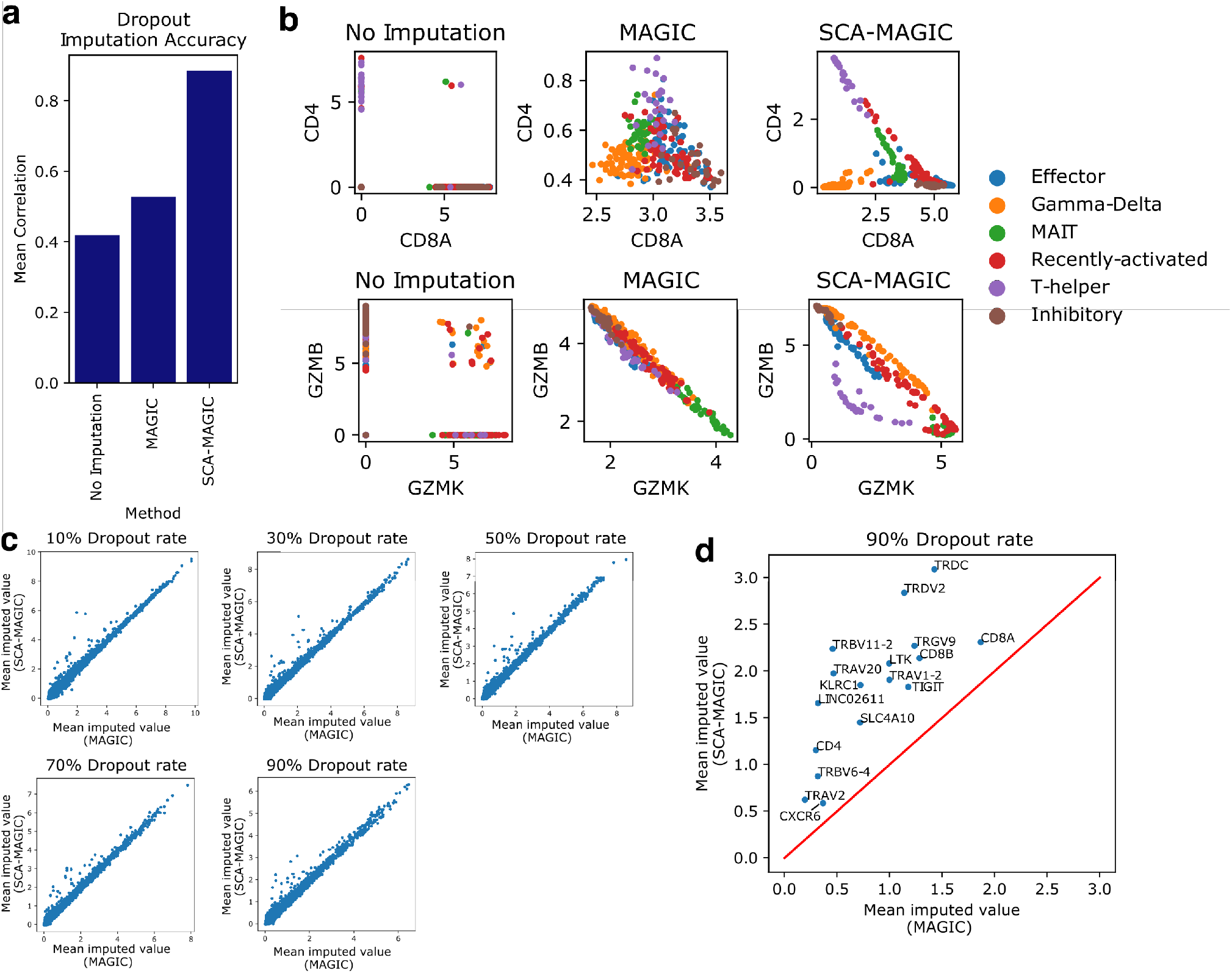
Imputation performance of MAGIC using PCA and using SCA (SCA-MAGIC). **a**: Recovery of marker genes on the 500-cell dataset analyzed in Figure 2. For each method, we measure the average correlation between the imputed marker genes and an indicator vector for the rare cells. Our version, SCA-MAGIC, achieves significantly higher correlation. **b**: Visualizing gene-gene relationships in cytotoxic T-cell data after imputation using MAGIC and SCA-MAGIC. SCA-MAGIC better recovers the inverse relationships between CD8 and CD4 and between Granzyme B and Granzyme K, with gamma-delta T-cells expressing neither CD8 nor CD4 and lower Granzyme expression for T-helper cells. **c**: Scatter plot showing mRNA dropout recovery in the cytotoxic T-cell dataset at various dropout rates. A fixed percentage of nonzero transcript measurements were set to zero, and the mean imputed values of these removed transcripts were assessed for each gene. While MAGIC and SCA-MAGIC perform similarly on most genes, SCA-MAGIC consistently outperforms MAGIC on a subset of them. **d**: A closer look at the genes where SCA-MAGIC significantly outperforms MAGIC in recovering dropouts at the 90% dropout rate; they include many key marker genes such as *CD8A, CD8B, CD4, TIGIT*, and *TRDV2*.

To assess recovery of gene-gene relationships, we ran MAGIC and SCA-MAGIC on the cytotoxic T-cells from Hagemann-Jensen et al. [19]. We find that SCA-MAGIC better recovers the complimentary relationship between CD8 expression and CD4 expression among alpha-beta T-cells, with T-helper cells having high CD4 levels and gamma-delta T-cells expressing neither surface receptor, consistent with the literature [23, 30] (Figure 5b, top). Both MAGIC and SCA-MAGIC report an inverse relationship between the expression levels of Granzyme B and Granzyme K. However, SCA-based MAGIC assigns the T-helper cells lower Granzyme B levels than the other populations. This is concordant with flow-cytometry results, which indicate that CD8+ T-cells are the primary secretors of granzyme B [31].

To compare dropout recovery on real data, we created low-coverage versions of the cytotoxic T-cell dataset by setting 10%, 30%, 50%, or 90% of the nonzero transcript counts to zero, and examined the values of these dropped-out transcripts after imputation. Higher imputed values indicate better transcript recovery. For most genes, SCA-MAGIC and MAGIC perform similarly (Figure 5c). However, SCA-MAGIC outperforms MAGIC on a small set of key marker genes, including *CD8A, CD8B, CD4, TIGIT, KLRC1*, and the gamma-delta marker genes *TRDV2* and *TRGV9* (Figure 5d). Since these genes define important T-cell subclasses, this improvement is consequential.

### SCA scales to large datasets

As improving technologies generate ever larger datasets, the computational tools used to analyze these datasets must scale accordingly. SCA meets this need with fast runtimes and modest memory overhead.

Asymptotically, SCA’s runtime and memory overhead are both linear in the size of the input dataset (Supplementary Note 2, Figure S1). To test this empirically, we measured runtime and peak memory usage for one iteration of SCA on data of varying sizes (Figure S1). We produced test datasets by taking random subsets of Patient 1’s T-cell data from Hao et al. [5] with varying numbers of cells and genes. For all tests, we use a neighborhood size of 15. As expected, we find SCA’s runtime and memory performance scale linearly with increasing numbers of cells. On a subset with 9,000 cells and over 20,000 genes, SCA takes 3 minutes and 15 seconds to run with a peak allocation of 561 MB. This is only slightly slower than PCA, which finishes in 2 minutes and 10 seconds and allocates 179MB. (ICA is somewhat more computationally demanding, requiring 266 seconds and allocating over 4GB). SCA’s linear scaling makes it tractable even on the very largest single-cell datasets; for example, on a mouse brain dataset from Saunders et al. with 939,489 cells and 20,658 genes [3], SCA runs in 5.5 hours and allocates a maximum of about 29GB, well within the range of most modern day computers. Sub-sampling techniques that preserve rare cell types, such as Hopper [13] and Geosketch [12], may be combined with SCA to enable fine-grained analyses of these massive datasets even on a laptop. We can also reduce the memory overhead further by processing the data in chunks containing a certain number of genes (Methods).

## Discussion

SCA offers an information-theoretic approach to measuring and compressing salient transcriptional signals in single-cell data, enabling downstream analyses at unprecedented resolution. By iteratively boosting the locality-specific signal of individual transcripts, SCA uncovers clinically-relevant immunological populations that are invisible to existing approaches.

A variety of approaches have arisen that are specifically tailored to the problem of rare cell type recovery. However, we find that these methods have limiting assumptions [9, 11] or rely on potentially inaccurate or ill-defined clustering procedures [10, 8] that limit performance. GiniClust [9] assumes that genes with high Gini index are the most important; however, we demonstrated that this is not always the case (e.g. the marker genes in the synthetic dataset were not marked as having high-Gini index by the algorithm). RaceID and CellSIUS both compute initial clusterings which are then individually refined. However, an accurate initial clustering may be difficult to obtain when cell types of interest are rare or subtly-defined, and cluster-based approaches are unsuitable when cells form more continuous transcriptional structures, such as developmental trajectories, which do not neatly partition [32]. FiRE [11] sidesteps the limitations of clustering by assigning each cell a rarity score according to its degree of isolation, but the notion of isolation in turn relies on a meaningful cell-to-cell distance metric, which is not readily derived. Hopper [13] reduces the data in the hopes of increasing the proportion of rare cells, but its approach requires a reliable distance metric, and requires getting rid of potentially valuable data. For the latter two methods, one might improve performance by using Euclidean distance in an SCA representation as a distance metric. Our work suggests that the right dimensionality reduction can enable recovery of even rare and subtly-defined populations.

SCA’s surprisal scores are similar in principle to the Inverse Document Frequency (IDF) transform, a normalization approach widely used in text processing and in some single-cell applications, whereby each feature (gene) is weighted by the logarithm of its inverse frequency [33]. Like SCA, IDF gives rarely-seen features more weight; however, it does not consider the locality-specific context of each feature measurement, so it lacks the statistical power to detect locally-enriched signals. By incorporating counts from local neighborhoods of each cell, SCA allows genes to have variable scores across the dataset, achieving high-magnitude scores where they are discriminative and near-zero scores where they are noise (Figure 1c). Our approach reflects true biology, where genes may be expressed sporadically across the entire dataset but mark informative distinctions only within a small subpopulation.

SCA is also conceptually similar to surprisal analysis [34], which compares observed data to a pre-computed balance state to identify meaningful deviations. Originally developed for thermodynamics, these methods have recently found use in *bulk* transcriptomic analysis of biological systems in flux, such as cancer cells undergoing epithelial-to-mesenchymal transition and carcinogenesis [35, 36, 37, 38]. For example, Gross et al. [35] perform singular value decomposition on a surprisal matrix derived from time series micro-arrays to identify bulk transcriptomic signatures that predict eventual malignancy. In their work, the surprisal of a transcript is defined by the negative logarithmic fold change of the transcript from its value in the balance state. We generalized this idea to single-cell data by treating each cell as a separate time point, and computing surprisals as negative log-fold changes between observed transcript counts and transcript expression means across all cells. However, we show this extended notion of surprisal is under-powered and inaccurate for single-cell data (Supplementary Note 5, figure S6), because individual transcript counts are themselves noisy. For example, on the cytotoxic T-cell dataset, this approach fails to separate CD8 from CD4 T-cells (Figure S6). SCA’s novel approach of testing expression in *neighborhoods* of cells instead of individual cells lends statistical power and limits the impact of noise and dropouts, especially in combination with the robust Wilcoxon test.

Data visualization, which features prominently in many single-cell pipelines [39, 20], differs from dimensionality reduction, on which we focus. Whereas visualization aims to produce a two-dimensional rendering of the data, dimensionality reduction produces a smaller, but still many-dimensional representation which is then analyzed further downstream. Thus, data visualization tools compliment dimensionality reduction rather than substitute for it. Indeed, visualizations are often built on dimensionality-reduced data; for example, UMAP plots in existing literature are often computed on PCA or ICA reductions [3, 40]. SCA complements existing visualization tools to facilitate exploratory analysis (Figure 4a, 3a, and S4).

SCA also combines well with *sketching* techniques, such as Geosketch [12] and Hopper [13], which generate subsamples of cells that retain transcriptional diversity; SCA can in principle be run on the subsamples from these methods to allow massive single-cell analysis on a laptop, thus democratizing single-cell analysis. In turn, these sketching techniques rely on a low-dimensional representation of cells, which SCA may provide. As motivation for the latter, we have shown that SCA is better at identifying rare cell types than these sketching techniques.

Although the process that generates the surprisal components is nonlinear, requiring nearest-neighbor graphs and Wilcoxon score computation, SCA’s output is a linear projection of its input. This places SCA firmly in the linear category, together with PCA and ICA; indeed, for fixed dimension *D*, the coordinate systems defined by SCA and by these methods are related by rotation in the original high-dimensional space. Intuitively, SCA changes the “perspective” from which the data is viewed. It is remarkable, then, that SCA’s reductions look so different in downstream analyses from those of PCA and ICA (e.g. Figure 3a). This is possible because high-dimensional space offers a far wider variety of perspectives than the three-dimensional space we often think in, giving linear methods more richness than they are usually credited for.

Dimensionality reduction addresses the underlying goal of nearly all single-cell analytic pipelines – to determine which cells are phenotypically similar to one another, or in mathematical terms, to derive a biologically meaningful metric between cells. If we could meet this goal perfectly, we could immediately obtain perfect clusterings of single-cell data (each cluster would be a connected component of the *k*-nearest neighbor graph), perform perfect batch correction (by integrating cells based on phenotypic similarity), and substantially improve trajectory inference (by connecting similar cells along a continuous path). Dimensionality reduction represents single-cell data in a lower-dimensional Euclidean space, which inherits natural metrics (e.g. the standard Euclidean distance). SCA provides an embedding where Euclidean distance better captures biological similarity, causing cells with similar phenotypes to cluster together.

### Software availability

SCA is implemented in Python, installed on PyPi for ease of use, and offers integration with scanpy, a popular scRNA processing framework. Links to the source code, installation instructions, and complete documentation are available at `sca.csail.mit.edu`.

## Methods

### Synthetic Data Experiments

The synthetic dataset analyzed in Figure 2 was generated using numpy [41] and scanpy [42]. Nonzero expression values are drawn randomly with replacement from the nonzero log-TPM expression values of the cytotoxic T-cell dataset from [19], which is analyzed in Figure 3 and described below.

The locations of the nonzero expression values are chosen to noisily divide the data into a “common” population of 475 cells and a “rare” population of 25 cells (5% of the total). 20 “marker” genes are expressed with frequency 90% in the rare population, and 15% in the common population. The remaining 1,000 “noise” genes are expressed uniformly at random, with each cell having a 20% chance of expressing each noise gene. For PCA representations, we compute 50 PCs using scanpy’s `pca` function [42], which in turn uses the implementation of `scikit-learn` [43], with default parameters. We performed ICA using the `scikit-learn` implementation of `fastICA` [43, 44]. We ran SCA with a neighborhood size of 15 for 1-4 iterations with 20 components, and an automatically-determined multiple testing correction factor of 301 (see below and Algorithm S1). In each representation, we computed the 15-nearest-neighbor graph using the Euclidean distance metric. The resulting neighborhood graphs were then used to generate the UMAP plots shown in Figure 2b, using `scanpy`’s implementation with default parameters [42, 20]. Silhouette scores were computed using the `silhouette score` method from `scikit-learn` [43, 21].

### T-cell data from Hao et al

We obtained transcriptomic data from the authors’ website at https://atlas.fredhutch.org/nygc/multimodal-pbmc/. The log-transformed count data was subset to T-cells using the authors’ annotations, yielding 73,259 T-cells and 20,729 genes across all 8 patients. We then split by patient into 8 donor-specific datasets. For each donor, we computed an SCA reduction using 50 components and 5 iterations, with a neighborhood size of 100 (a larger neighborhood size than that used for the cytotoxic T-cell data is appropriate, because the input data is larger). 50 principal components were computed using scanpy’s `pca` function, and 50 independent components were computed with the `FastICA` implementation provided by `scikit-learn` [44, 43]. To ensure convergence of `FastICA`, we raised the maximum number of iterations to 500 from the default of 200. We ran scVI with default parameters (learning rate 0.001, 400 warmup epochs for KL divergence term), and a 50-dimensional latent embedding space to match the dimensionality of the PCA, ICA, and SCA embeddings. We computed 15-nearest-neighbor graphs in each representation using the Euclidean distance metric, then ran UMAP [20] and Leiden clustering [27] on the resulting neighborhood graphs. For Leiden clustering, we use the default resolution of 1.0.

We used the following procedure to identify a subset *S* of clusters achieving high F1 score for each target population *P*, from a set of *T* clusters *c*_1_, …, *c*_*T*_.

1. Rank all clusters by the degree of overlap with *P*, i.e. by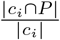. Ties may be handled arbitrarily. Assume without loss of generality that the indexing *c*_1_, …, *c*_*T*_ ranks the clusters in this way. Initialize *S* to be empty.
2. for *i* in 1,2,…,*T* :
  a. Measure the F1 score of *S* ⋃ *c*_*i*_ with respect to *P*.
  b. If this F1 score is higher than that of *S*, add *c*_*i*_ to *S*. Otherwise, stop and return the current F1 score of *S* with respect to *P*.
3. Return the F1 score of *S* (if not already returned above).

If the target population is a union of clusters, this procedure is guaranteed to find it; otherwise, it finds a set of clusters which approximates *P*.

### Cytotoxic T-cell Population Discovery

We extracted all cytotoxic T-cells from the SS3 dataset in [19] using the authors’ cell type annotations, obtaining 307 cells in total. For PCA, we used scanpy’s `pca` function with 20 components. For ICA, we used the `FastICA` function of `sklearn` [43, 44], again with 20 components. For SCA, we ran five iterations with 20 components each, starting with the PCA representation. We ran scVI with default parameters on the top 4000 most variable genes (we observed little difference in performance when running on all genes). 15-nearest neighbors graphs were computed in each representation using Euclidean distance, and the results were used to generate the UMAP plots in Figure 3. Leiden clusters in each representation were computed with the default resolution of 1.0. Matrix plots to show expression of key marker genes were generated using scanpy’s `matrixplot` function.

### Imputation using MAGIC

To create SCA-MAGIC, we use the `graphtools` package to build a diffusion operator based on a 20-dimensional SCA reduction, with default parameters inherited from MAGIC (*knn=5, knn max=15, decay=1, thresh=0*.*0001*). We use the same parameters to construct an analogous operator from a 20-dimensional PCA embedding. We then build `MAGIC` instances from these two operators, with time parameter *t=5*, and compare performance on various datasets.

To generate artificial dropouts in the cytotoxic T-cell data, we replaced a random subset of the pooled nonzero transcript measurements with zeros, comprising either 10%, 30%, 50%, or 90% of the total nonzero measurements. After performing imputation, we re-examined the transcripts that had been eliminated, and checked whether they had been restored. High imputed values indicate well-restored transcripts.

### Details of the SCA Algorithm

#### Surprisal score matrix computation (Algorithm 1)

Given the input data *X* with *N* cells and *M* genes, a target dimensionality *D*, and a neighborhood size *k*, we first compute a *D*-dimensional PCA reduction of *X*. Using Euclidean distance in this PCA space, we compute for each cell *c* a neighborhood *N*_*k*_(*c*) containing the *k* nearest cells. Alternatively, the user may specify neighborhoods manually as lists of indices.

For each gene *g* and cell *c*, we then assess the significance of *g*’s expression in *N*_*k*_(*c*) as compared to its global expression. Under the null hypothesis, where *g* is randomly expressed, the local distribution *N*_*k*_(*c*) should be similar to the global expression. Using a Wilcoxon rank-sum test, we obtain a two-sided *p*-value *p*_*c,g*_ representing the probability of the observed difference under this null hypothesis. We also offer two alternative *p*-values based on different models: a t-test, and a binomial test using only the binarized counts. We strongly recommend the Wilcoxon model for its flexibility to a wide range of data distributions, and robustness to different pre-processing protocols.

Small *p*-values indicate very unlikely events under the null hypothesis, leading to high surprisal. However, since tens of thousands of genes are often measured for each cell, we would expect *p*_*c,g*_ to be very low for some cell-gene combinations even in the absence of true biological signal. We therefore adjust for multiple-testing within each cell using a family-wise error rate correction. If we assume that genes are uncorrelated, this correction takes the form

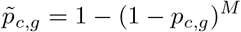

where *M* is the number of genes. The corrected value 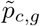 represents the probability that *any* gene has the observed deviation from the null distribution in *c*’s neighborhood. However, in real single-cell data, genes are often highly correlated, so the effective number of independent features is far fewer than *M*. As detailed in Supplementary Note 4 and Algorithm S1, we can identify a reasonable exponent *N*_*t*_ by sampling many random sets of *k* cells from *X*, computing *p*-values from these random neighborhoods, and observing the distribution of these *p*-values. This provides a background model for contextualizing the *p*_*c,g*_ values computed from the actual locally derived *k*-nearest neighborhoods, and leads to the correction

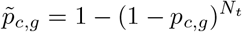

where *N*_*t*_ is often far less than *M*. When *N*_*t*_ is computed as in Supplementary Note 4, SCA does not produce erroneous clusters on negative control datasets which lack intrinsic structure, and randomly-generated neighborhoods yield scores clustered around zero (Supplementary Note 3, Supplementary Figure S3). SCA computes *N*_*t*_ in this way by default; however, users may also manually define *N*_*t*_ to adjust the balance between sensitivity and specificity.

We next convert the corrected *p*-values 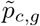 into surprisal scores *S*(*c, g*). Shannon [18] defines the *surprisal* or *self-information* of an event with probability *p* as *-* log(*p*). Intuitively, less probable events are more informative when they occur. For a given cell *c* and gene *g*, 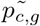 is the probability of the event that that one of *c*’s genes has a local distribution at least as extreme as the observed distribution of *g*, under the null hypothesis of random gene expression. Thus, 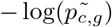 is the surprisal of this event, and defines the magnitude of *S*(*c, g*).

To distinguish over-expression from under-expression, we give *S*(*c, g*) a positive sign if *g* is over-expressed in *c*’s neighborhood and a negative sign if it is under-expressed. Under the Wilcoxon model, over- or under-expression is determined by the sum of the ranks of *g*’s values in the *k*-neighborhood of *c* among all values *g* takes, which we denote ranksum(*g, N*_*k*_(*c*)). Under the null hypothesis, this quantity follows a normal distribution with mean 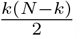. Thus, we obtain

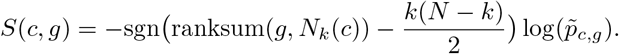

### Computing surprisal components (Algorithm 2)

From the surprisal scores *S*(*c, g*), SCA next seeks to generate an informative linear combination of genes. For a given combination defined by

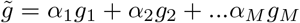

we say that 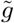 has *loadings α*_1_, …, *α*_*n*_. For a fixed cell *c*, we formulate the surprisal score of 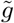 as

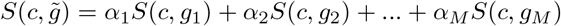

We then define the overall overall surprisal score of 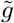 by taking the norm over all cells:

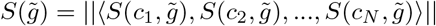

or, in matrix notation,

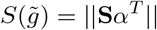

where **S** denotes the surprisal score matrix and *α* = *hα*_1_, …, *α*_*M*_ *i*.

We now seek the metagene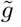, defined by the loadings *α*_1_, …, *α*_*N*_, that maximizes 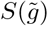. Since we can achieve arbitrarily large values of 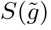 by scaling the coefficients, we constrain the loading coefficients to have norm 1, that is:

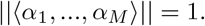

It is a standard linear algebra result that this maximum is realized by the leading right-eigenvector of **S** (proof in Supplementary Note 1 and [45]). Thus, the first surprisal component loading vector is simply the first right eigenvector of **S**, which we denote **v**_**1**_. To obtain additional surprisal components, we repeat the optimization with the constraint

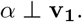

It is straightforward to see (Supplementary Note 1) that this yields the second principal component loading vector **v**_**2**_ of **S**. Continuing, we see that the loading vectors for surprisal components are simply the right eigenvectors of **S**.

SCA next computes the values of the first *D* surprisal components over the *input* data (not the surprisal scores). That is,

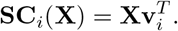

Note that the surprisal components are linear functions of the input data, despite the nonlinear construction of **S**. Although the loadings are computed on **S**, the values of the components are computed by applying these loadings to **X**.

If desired, we can now use the resulting *D*-dimensional representation of *X* to compute a Euclidean *k*-nearest neighbor graph, compute a new surprisal score matrix from these neighborhoods, and perform SCA on this new matrix to produce another *D*-dimensional representation of *X*. This can be repeated arbitrarily many times, and often improves performance up to 3-4 iterations (Figure 3f, Figure S2).

### Time and memory optimizations

Computing the surprisal scores *S*(*c, g*) for all cells *c* and genes *g* requires *NM* Wilcoxon rank-sum tests. However, we can rapidly produce all of the rank-sum statistics with minimal memory overhead as follows:

1. Divide the genes into chunks of a user-specified size *C*, depending on memory constraints (default 1000). Let *G*_1_ = *{g*_1_, …, *g*_*C*_*}, G*2 = *{g*_*C*+1_,, *g*_2*C*_*}*, and so on.
2. For each gene chunk *G*_*i*_:
  a. Subset *X* to genes in *G*_*i*_, obtaining a reduced dataset *X*_*i*_
  b. rank each column of *X*_*i*_ to obtain a rank matrix *R*_*i*_
  c. Multiply the adjacency matrix *A* with *R*_*i*_, yielding a rank-sum matrix over neighborhoods, denoted *RS*_*i*_, overwriting *R*_*i*_
  d. Convert these rank-sums into *p*-values under to the null model, overwriting *RS*_*i*_ with a p-value matrix *P*_*i*_
  e. Convert these *p*-values into surprisal scores, as described in Algorithm 1, overwriting *P*_*i*_ with surprisal scores *S*_*i*_.
  f. Sparsify *S*_*i*_ and record it. (*S*_*i*_ is frequently quite sparse).
3. Concatenate the matrices *S*_*i*_ horizontally to obtain the surprisal score matrix *S*.

Using this approach, we only need to compute ranks for each gene once, and we avoid storing dense matrices of size larger than *N × C*. Since *A* has at most *k* nonzero elements per row, the sparse matrix multiplication in step 2c requires only *O*(*kNC*) time. The remaining steps are easily accomplished with vectorized functions from `scipy` [46] and `numpy` [41].

With these improvements, SCA is nearly as fast as ICA and PCA, and uses significantly less memory than ICA. For example, running SCA on Patient 1 from the Hao et al. dataset (9,000 cells and 20,000 genes) finishes in about 5 minutes and allocates about 500MB of RAM (Figure S1). We include more in-depth time and memory benchmarks in Supplementary Note 2 and Figure S1.

## Acknowledgements

The authors are grateful to Ashwin Narayan, Brian Hie, Hyunghoon Cho, Sarah Nyquist, Rohit Singh, Ellen Zhong, and other members of the Berger lab for their continual support and helpful feedback. We also thank Josh Peters, Bryan Bryson, and Cheng-Zhong Zhang for helpful conversations on SCA’s applications and valuable suggestions for biological applications. This work was supported by the National Institutes of Health U01CA250554 [to B.B.].

## Algorithm 1: InfoScore

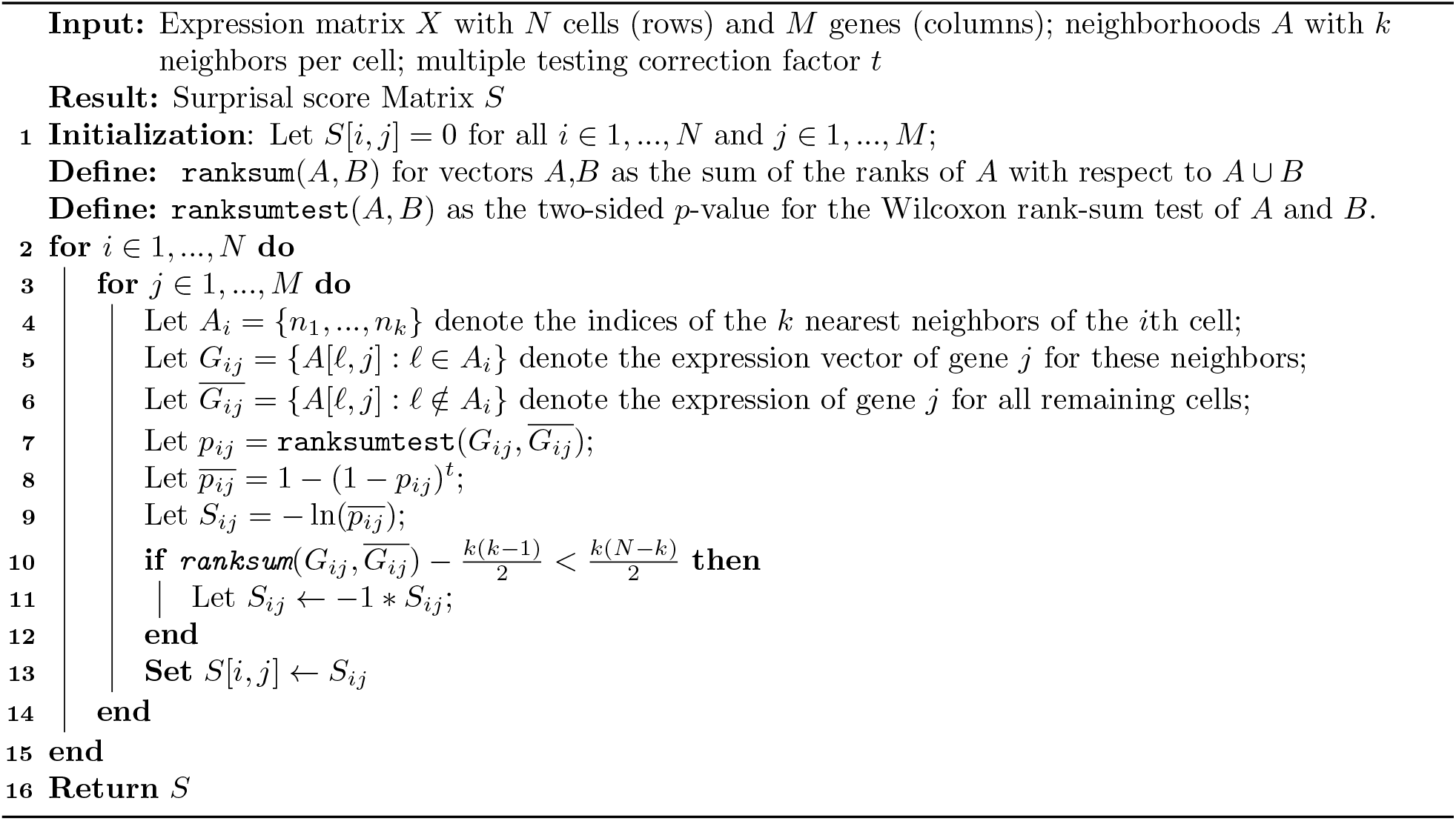

## Algorithm 2: SCA

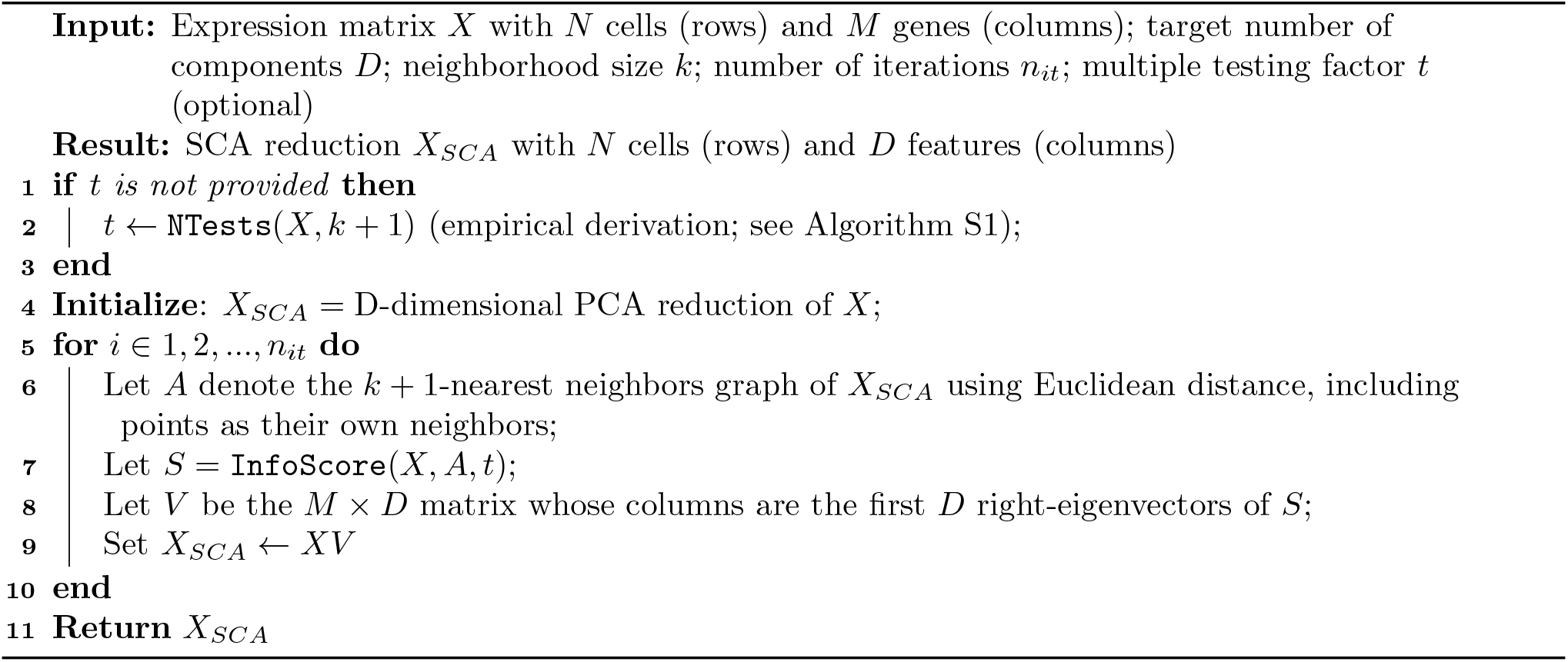

