## Supplementary Information for "Recovering Single-cell Heterogeneity Through Information-based Dimensionality Reduction"

November 12, 2021

#### Supplementary Note 1: Mathematical Formalization

The main text uses the following lemma to justify SCA:

**Lemma 1.** *If $X$ is an $n \times m$ matrix and $\alpha$ is an $m$-dimensional vector with unit $L_2$ norm, then $||X\alpha^T||$ is maximized when $\alpha$ is a principal right-eigenvector of $X$.*

The following proof is adapted from [1]:

*Proof.* Note that

$$||X\alpha^T||^2 = \langle X\alpha^T, X\alpha^T \rangle = \langle X^T X\alpha^T, \alpha^T \rangle.$$

Since $X^T X$ is symmetric, it admits a set of orthonormal eigenvectors $v_1, ..., v_k$ with $X^T X v_k = \mu_k^2 v_k$ and positive eigenvalues $\mu_1^2 \geq \mu_2^2 \geq ... \geq \mu_k^2$. Now, express $\alpha$ in the eigenbasis of $X^T X$:

$$\alpha = \sum_{k=1}^{m} a_k v_k.$$

Note that since the vectors $v_i$ are orthonormal and $\alpha$ is a unit vector, we must have

$$||\langle a_1, ..., a_k \rangle|| = 1.$$

Now,

$$\langle X^T X\alpha, \alpha\rangle = \langle \sum_{k=1}^{m} a_k\, X^T X v_k, \sum_{k=1}^{m} a_k v_k\rangle = \sum_{j,k=1}^{m} a_j a_k \langle X^T X v_j, v_k\rangle \tag{1}$$

$$= \sum_{j,k=1}^{m} a_j a_k \langle \mu_k^2\, v_k, v_j\rangle = \sum_{j,k=1}^{m} a_j a_k \mu_k^2\, \langle\, v_k, v_j\rangle \tag{2}$$

$$= \sum_{k=1}^{n} a_k^2\, \mu_k^2. \tag{3}$$

Since $||\langle a_1, ..., a_k\rangle|| = 1$, this is maximized by setting $a_1 = 1$ and all other $a_i = 0$, yielding $\alpha = v_1$. This completes the proof □

In particular, this validates the use of right-eigenvectors of $I$ as components with high surprisal.

We now seek additional maximally-informative components orthogonal to the first right-eigenvector of $I$. This is equivalent to constraining $a_1 = 0$ in (3); in this case, the maximum is clearly attained by setting $a_2 = 1$ and all other $x_i = 0$, resulting in $\mathbf{a} = v_2$. Continuing, we see that the right eigenvectors yield an orthonormal basis which is maximally informative in the sense outlined in the main text.

#### Supplementary Note 2: Time and Memory Performance

In light of rapidly increasing single-cell dataset sizes, it is vital that emerging methods scale to large datasets. To see how SCA scales, we ran SCA for a single iteration on various-sized subsets of the T-cell from Patient 1 from Hao et al. [2], containing 9,969 T-cells and 20,739 genes. To assess scaling with increasing numbers of cells, we measure time and memory performance on slices of Patient 1's T-cell data with all genes and random cell subsets of size 1000, 2000, 3000,...,9000. Similarly, to assess scaling with increasing numbers of genes, we assess performance on slices with all cells and random gene subsets of size 4000-20000, in increments of 4000. For each slice, we run three replicates to ensure robust measurements. All benchmarks were measured on an Intel Xeon Gold 6130 CPU (2.10GHz, 768 GB of RAM).

The results of this analysis, shown in Figure S1a, show that SCA's runtime and memory footprint are linear the number of cells. As described in the Methods, SCA processes data in constant-sized chunks of genes to minimize memory allocation. For these experiments, we group genes into chunks of size 1000; consequently, the memory allocation does not depend strongly on the total number of genes. SCA is necessarily slower than PCA, because it runs a singular-value decomposition analogous to PCA's on the surprisal score matrix.

For further comparison, we also performed time benchmarks for CellSIUS [3], GiniClust [4], FiRE [5], and RaceID [6] on the entire Patient 1 data from Hao (9,969 T-cells and 20,739 genes; Figure S1b). We found that all of these methods have comparable runtime with the exception of RaceID, which takes nearly two hours to run.

#### Supplementary Note 3: Performance on randomized data

A major concern with any signal-boosting approach is false positives, i.e. finding signal when none is present. For SCA, we must ensure that truly structure-less data remains so in the SCA representation.

We construct a random Gaussian dataset (i.e. each transcript is drawn at random from a standard Normal distribution) with 1,000 cells and 10,000 genes. As expected, UMAP plots derived from 20-dimensional SCA representations do not show any significant structure (Figure S3a). As a more realistic scenario, we randomly permuted each gene in the cytotoxic T-cell dataset from [7], eliminating gene-gene correlations (Figure S3b). On this data, the leading PC aligns with the highest-variance gene, in this case granulysin, so that the PCA representation loosely separates cells that express granulysin from cells that do not. SCA boosts this signal to produce a stronger separation, but does not otherwise cluster the data.

#### Supplementary Note 4: Automatic determination of the multiple-testing correction factor

Transcriptional datasets often profile tens of thousands of genes, but correlations among the genes mean that the data has significantly fewer degrees of freedom. Consequently, the statistical tests performed to produce the Information score matrix may be highly dependent. This affects the degree of multiple testing correction these scores should undergo. For example, if the genes partition into two highly-correlated modules, then each cell really only performs two tests, for the over- or under-expression of these modules. We therefore devised an empirical method to determine the appropriate multiple testing correction factor (Algorithm S1). The basic idea is to compute minimum $p$-values across all genes over many random collections of cells, from which the appropriate exponent can be determined.

More formally, we generate $N$ random neighborhoods from a dataset with $M$ genes, yielding an $N \times M$ wilcoxon $p$-values. Let $p_{ij}$ denote the $p$-value for the $j$th transcript in the $i$th random neighborhood. Now let

$$\tilde{p}_i = \min(\{p_{ij} : 1 \leq j \leq M\}).$$

Note that, by construction, for all $P \in [0, 1]$, the $p_{ij}$ values are uniform in expectation, that is:

$$\mathbb{P}(p_{ij} < P) = P.$$

If all of the genes were independent of each other, then we would expect the $\tilde{p}_i$s to have a distrution whose CDF is

$$\mathbb{P}(\tilde{p} < P) = 1 - (1 - P)^M$$

However, dependence among genes may reduce the exponent on the right hand side. To determine the correct exponent $t$, we empirically measure this CDF for our observations. Assuming without loss of generality that

$\tilde{p}_1 \leq \tilde{p}_2 \leq ... \leq \tilde{p}_N$, we have an empirical CDF satisfying:

$$\mathbb{P}_{emp}(\tilde{p} < p_i) = \frac{i}{N}.$$

Comparing this to the previous equation, we obtain a regression problem:

$$\frac{i}{N} \sim 1 - (1 - \tilde{p}_i)^t.$$

After some algebra, this becomes

$$\log(1 - \frac{i}{N}) \sim t \log(1 - \tilde{p}_i).$$

In real datasets, we indeed observe that the relationship between $\log(1 - \frac{i}{N})$ and $\log(1 - \tilde{p}_i)$ is approximately linear. Thus, a reasonable choice for $t$ is the slope of the regression line between $\log(1 - \frac{i}{N})$ and $\log(1 - \tilde{p}_i)$ for the computed $\tilde{p}_i$ (assumed to be indexed in loosely increasing order). This regression line necessarily passes through zero, so there is no need to fit an intercept. The $t$ output by Algorithm S1 is the mean-squared error estimate for this slope.

#### Supplementary Note 5: Single-cell surprisal analysis

Remacle et al. [8] and Gross et al. [9] use surprisal analysis on time series micro-array data to characterize the drivers of early carcinogenesis. Their approach models the observed expression as deviation from a balance state, and uses SVD to identify transcriptional programs that describe this deviation. The authors do not generalize this approach to work on single-cell data, so it is not clear how it could be applied to reduce the dimensionality of these datasets.

To see whether this approach could work in the single-cell domain, we propose an extension: rather than considering several bulk samples from different time points, treat each individual cell as its own sample, and treat them as these authors treat time points. The “maximal energy” balance state is characterized by uniform random expression of all genes, representing no constraints. Following [9], we then define the surprisal of a transcript as the log ratio between its observed expression in a cell and its mean expression over all cells. We then perform SVD on the resulting surprisal matrix to extract a smaller number of transcriptomic programs that define cellular behavior, yielding a lower-dimensional representation.

We applied this to the cytotoxic T-cell data from [7], producing a 20-dimensional representation (the same dimensionality as the SCA, ICA, and PCA representations produced in the main text). We find that key marker genes do not neatly separate in the resulting data; for example, CD8 T-cells mix with CD4 T-cells in the downstream UMAP plot, and F1 score analyses show that other key populations do not cleanly separate (Figure S6). We suspect that this approach does not have the statistical power to detect deviation from equilibrium in individual cells, due to the higher noise compared to bulk data. SCA boosts statistical power by considering *neighborhoods* of cells, giving context to each measurement, and allowing separation of these key populations.

---

**Algorithm S1:** NTests

**Input:** Expression matrix $X$ with $N$ cells (rows) and $M$ genes (columns); neighborhood size $k$; number of trials $T$

**Result:** Multiple-testing correction factor $t$

;
Let $A$ be a $T \times k$ matrix containing $T$ random sets of $k$ neighbor indices drawn from $1, ..., N$;
Let $S = \texttt{InfoScore}(X, A, t = 1)$;
Let $P$ be a $T \times k$ matrix defined by $P_{ij} = -\exp(|S_{ij}|)$ (matrix of $p$-values when neighborhoods are randomly chosen);
Let $Q$ be a $T$-dimensional vector of the row minima of $P$ ;
Sort $Q$ from smallest to largest;
Let $Y = \langle \frac{1}{T}, \frac{2}{T}, ..., \frac{T-1}{T}, 1 \rangle$;
Set $Y \leftarrow -\ln(1 - Y)$, element-wise;
Set $X \leftarrow -\ln(1 - Q)$, element-wise;
Let $t = \frac{\text{mean}(\{x_i y_i : i \in 1,..,T\})}{\text{mean}(\{x_i^2 : i \in 1,....,T\})}$ denote the slope of the least-squares regression line between $X$ and $Y$.;
Set $t \leftarrow \lceil t \rceil$ (ensuring it is an integer);
**Return** $t$

---

### Supplementary Figures

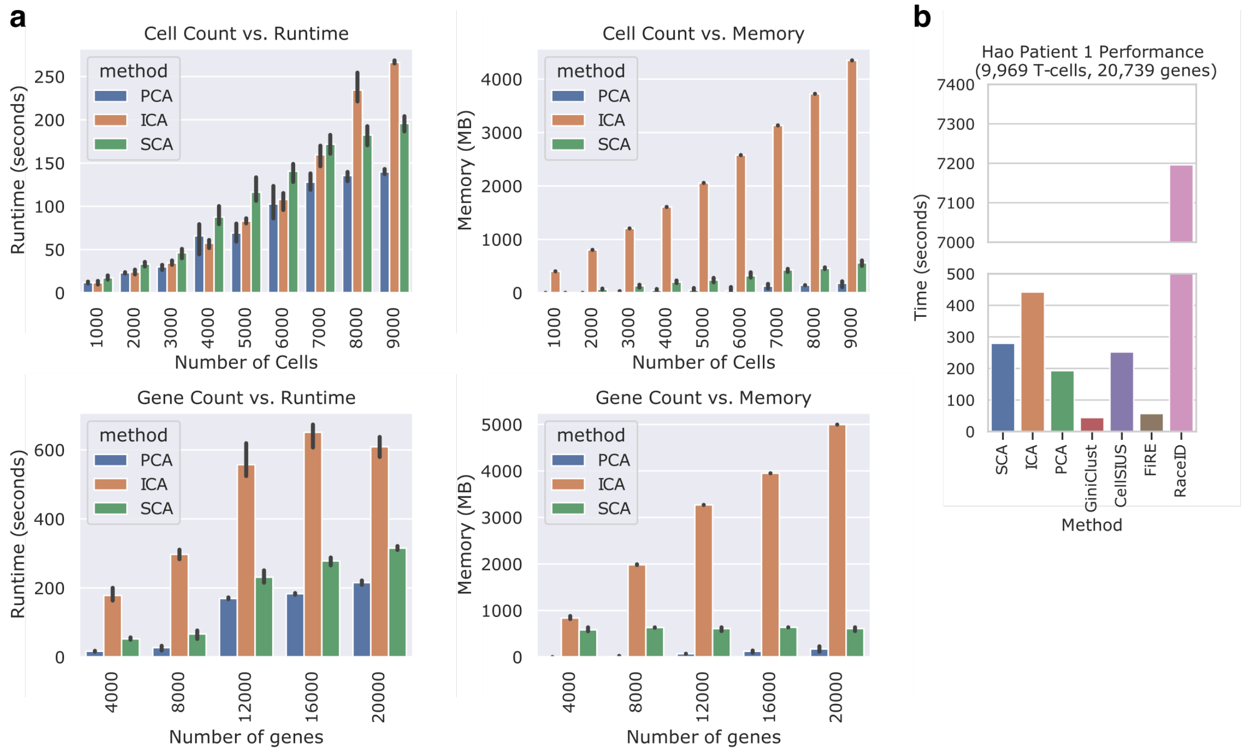


Figure S1: Time (left) and memory (right) benchmarks of SCA compared with PCA and ICA. The input dataset has 9,969 T-cells and 20,739 genes from Patient 1 in Hao [2]. In the top row, we assess scaling with increased cell numbers by taking random cell subsets of various sizes and measuring the time required to generate 50-dimensional reductions using SCA, PCA, or ICA. In the bottom row, we perform the same analysis on random gene subsets of varying sizes. Each bar represents three trials with different random subsets. (**c**) Runtime comparison of SCA against existing methods for rare cell type recovery on the full patient 1 dataset. RaceID takes by far the longest, requiring approximately 2 hours to run (note the broken axis).

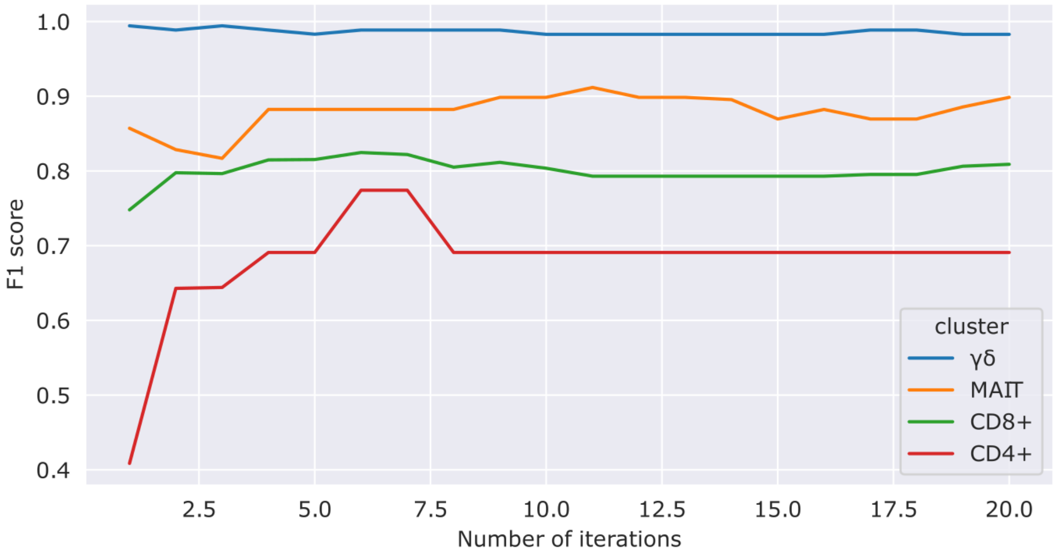


Figure S2: F1-score performance of SCA on the cytotoxic T-cell data from [7] with up to 20 iterations. Performance remains stable even after many iterations.

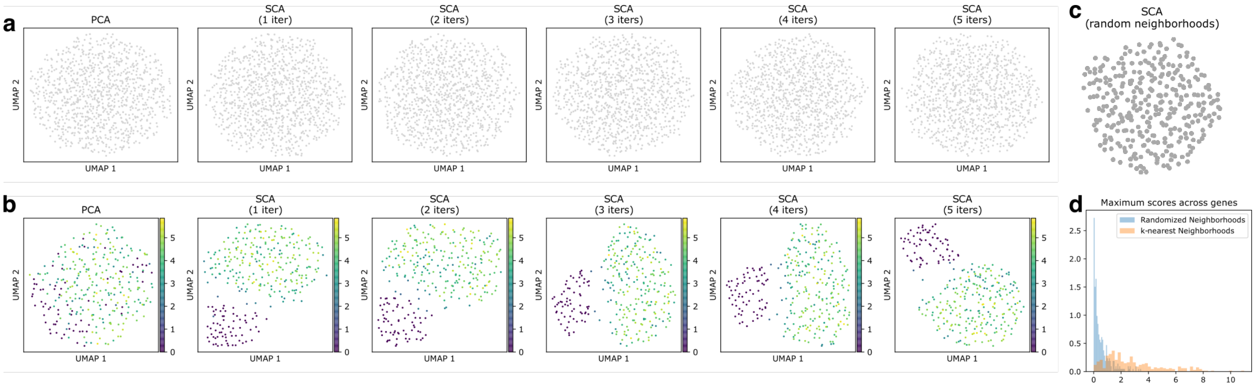


Figure S3: UMAP embeddings derived from SCA representations of "negative control" data without inter-feature relationships. We expect such data to have little structure, since most structure arises from correlations among many features. **a**: UMAP plots derived from SCA reductions of random Gaussian data with 1000 observations of 10,000 features. SCA does not reveal any structure even after 5 iterations. **b**: Reductions of scrambled cytotoxic T-cell data from [7]. Each gene was randomly permuted to eliminate gene-gene relationships. The plots are colored by granulysin expression. Since granulysin has high variance, the leading PC aligns with it, causing separation in the PCA representation which is amplified by SCA. **c**: UMAP plot derived from 1 iteration of SCA on the cytotoxic T-cell data, but where scores are computed on random neighborhoods instead of k-nearest neighborhoods. As expected, no obvious structure emerges **d**: Histogram of maximum magnitude of scores attained by each gene in any cell when neighborhoods are computer randomly (blue) or via k-nearest neighbors in PCA space (orange). When random neighborhoods are input, most scores are close to zero. When k-neighborhoods are used, the scores increase, reflecting additional information attained about the dataset's local structure.

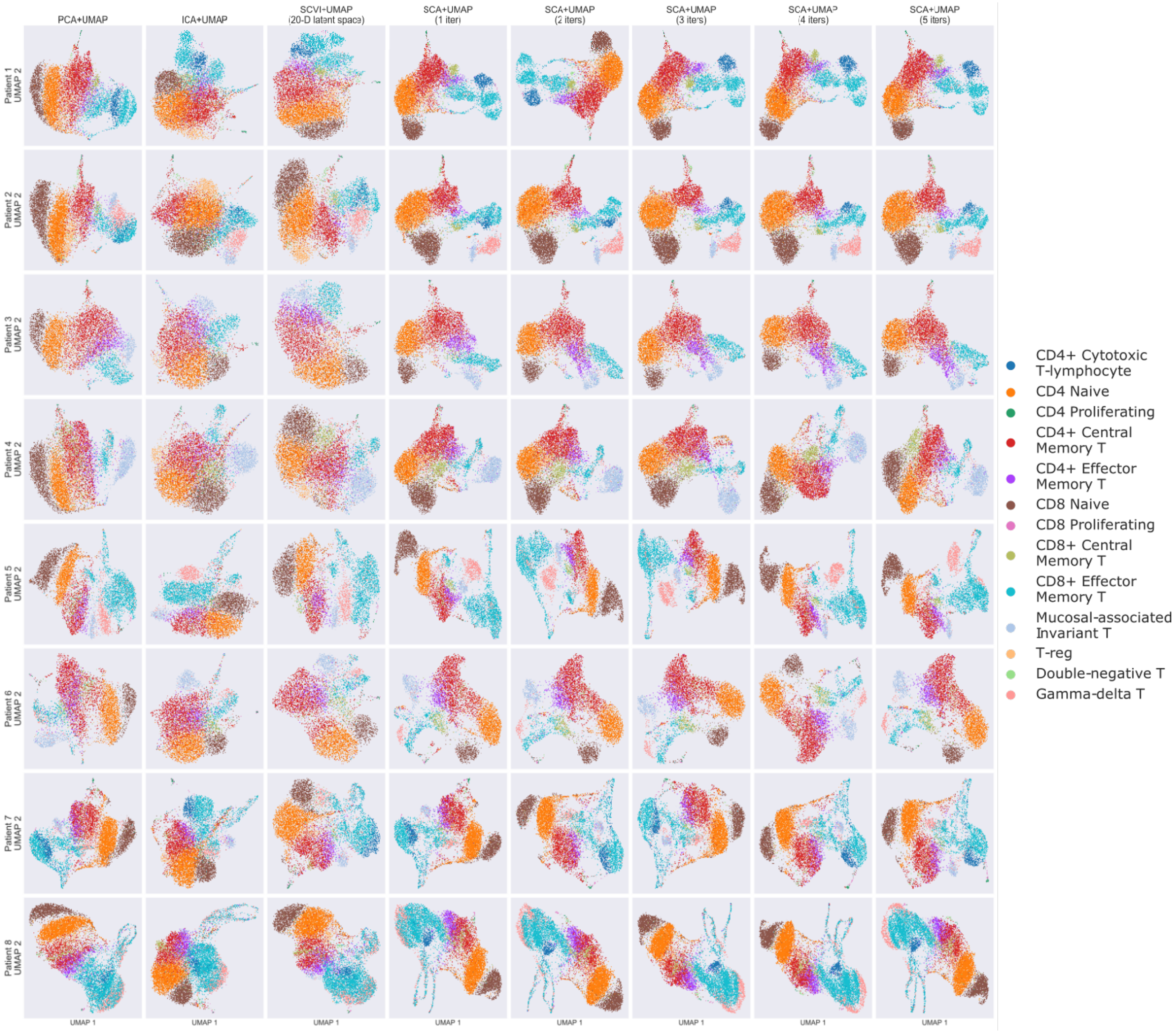


Figure S4: UMAP plots of cytotoxic T-cells for all patients in Hao et al., computed using PCA, ICA, scVI, or SCA with 1-5 iterations.

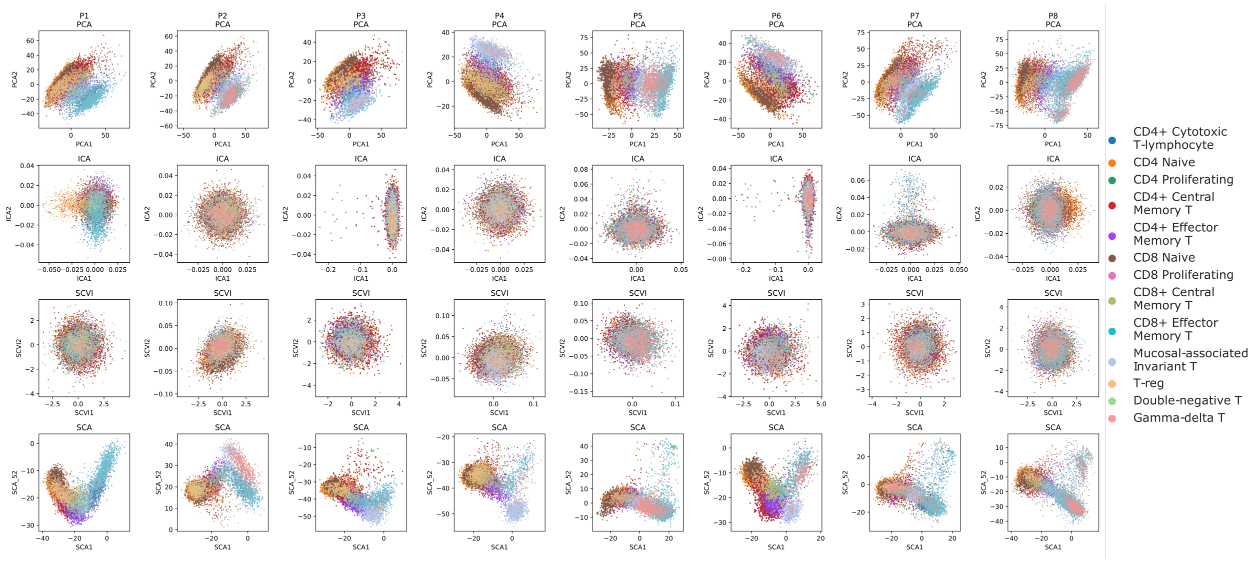


Figure S5: Component plots of cytotoxic T-cells for all patients in Hao et al., computed using PCA, ICA, scVI, or SCA with 1-5 iterations. For each reduction, the first two dimensions are plotted against each other. For SCA, we perform 5 iterations.

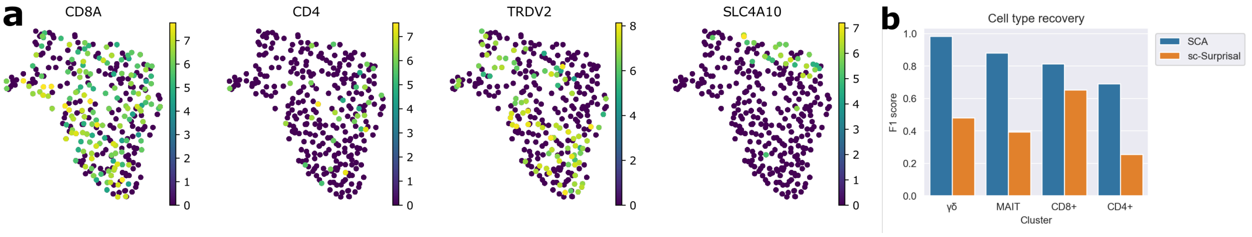


Figure S6: **a**: UMAP plots of the cytotoxic T-cell data after pre-processing using an extension of surprisal analysis used previously for only bulk transcriptomic time-series data [9]; in particular, we treat each cell as its own sample and consider it as a "time point" in the bulk approach. The surprisal of each transcript measurement is defined as the negative logarithm of its expression divided by its mean expression over all cells. We then perform 20-component SVD on the resulting matrix to produce a lower-dimensional embedding, which we input to UMAP. Key marker genes, like CD4, CD8, and TRDV2, do not separate well in the resulting embedding.**b**: F1 score analysis as in Figures 3 and 4. Leiden clusters downstream of SCA better capture known immunological subtypes.
